# Sex specificity of inhibitory gating deficits in individuals with high autistic traits

**DOI:** 10.64898/2026.08.11.744112

**Authors:** Mao Wen, Yufan Chen, Tianyi Gu, Binyuan Su, Pengmin Qin

**Author notes:** Corresponding Authors: Pengmin Qin, PhD and Binyuan Su, PhD, Pengmin Qin and Binyuan Su should be considered joint corresponding authors, with Pengmin Qin serving as the principal contact for all editorial correspondence.

## Abstract

Empirical evidence from traditional inhibitory control tasks regarding inhibitory deficits in high autistic traits has been mixed, indicating that this issue remains controversial. Although sex differences are widely documented in autistic cognitive profiles, their role in inhibitory gating mechanisms remains underexplored. Given that the expression of inhibitory gating deficits may be modulated by the social versus non-social nature of stimuli, and no prior study has investigated this topic by integrating both sex differences and stimulus domain, we addressed these two questions with the attribute amnesia paradigm. We manipulated stimulus type (non-social vs. social). In Experiment 1, participants performed a location task with animal drawings as targets and were unexpectedly asked to report animal identity on a surprise trial. High autistic trait females showed significantly higher accuracy on the surprise trial than all other groups, reflecting a failure to actively filter out task-irrelevant non-social information, that is, a reduced inhibitory gating efficiency. In Experiment 2, using face stimuli and a self-vs. other-face design, this gating deficit was no longer expressed: all groups performed at chance levels on the identity judgment, regardless of autistic trait level, sex, or face type. This dissociation aligns with a dual-mechanism framework: the inhibitory gating deficit in high autistic trait females is specific to non-social stimuli and masked by camouflaging for social ones. This study demonstrates that the inhibitory gating deficit in high autistic trait females is not a global impairment but rather a stimulus-dependent one, highlighting the need to consider sex and stimulus type.

## Introduction

The autism trait continuum framework, proposed by Baron-Cohen (1995), posits that autistic traits are not confined to clinically diagnosed autism spectrum disorder (ASD), but are distributed continuously across both clinical and general populations, with group differences lying primarily in the quantitative severity of traits (Baron-Cohen et al., 2001). Individuals with high autistic traits share core cognitive processing characteristics with those with ASD, including heightened attention to detail, superior rote memory, and difficulties in top-down attentional control (Hendry et al., 2020; Kunihira et al., 2006; Shah & Frith, 1993). Two dominant cognitive accounts have been proposed to explain these patterns. First, the weak central coherence account emphasizes a locally oriented processing style, marked by enhanced local feature processing but impaired global integration of information (Happé & Frith, 2006; Shah & Frith, 1983, 1993). Second, the enhanced perceptual functioning model further suggests that superior low-level perceptual operation in autism may also render individuals more vulnerable to interference from task-irrelevant stimuli (Lavie & Cox, 1997; Mottron et al., 2006). Together, these accounts point to a potential impairment in inhibitory working memory gating—the mechanism that actively blocks irrelevant sensory information from entering the capacity-limited working memory system.

However, behavioural evidence for this inhibitory deficit has remained inconsistent when tested with traditional paradigms such as the Simon task and Flanker task (Jones et al., 2021; Montgomery et al., 2021). A critical limitation of these tasks is that they mainly measure conflict resolution and behavioural inhibition at the response selection stage, and cannot isolate the earlier perceptual-encoding stage where attentional gating operates. Given that ASD-related filtering deficits are hypothesised to originate specifically at this early gating stage, conventional paradigms fail to capture the core impairment accurately.The attribute amnesia (AA) paradigm provides a precise approach to address this gap by directly dissociating attentional selection from working memory encoding. In a classic AA procedure, participants complete an ongoing task based on one stimulus attribute (e.g., location), and are unexpectedly asked to report another task-irrelevant attribute (e.g., identity) on a single surprise trial. The reliably lower accuracy on the surprise trial relative to subsequent informed control trials reflects active inhibitory gating: the brain actively suppresses attended but task-irrelevant information from entering working memory (Chen & Wyble, 2015a; Liu et al., 2025). Unlike traditional inhibitory tasks, the AA paradigm indexes working memory input gating in a relatively pure form, free from confounds of response conflict or motor inhibition. For individuals with high autistic traits, if their top-down inhibitory control over attentional inputs is weakened, they should show better memory for task-irrelevant attributes on the surprise trial (i.e., a reduced AA effect), making this paradigm ideal for testing the early inhibitory gating deficit hypothesis in autistic traits.

In the context of the AA paradigm, two factors may critically modulate the expression of this gating effect and thus need to be taken into account: sex differences and stimulus sociality. Regarding sex differences, accumulating evidence reveals substantial sex-related heterogeneity in ASD cognitive functioning and neurodevelopment. In the domain of executive function closely tied to inhibitory control, females with ASD show significantly more pronounced response inhibition impairments relative to typically developing females, whereas such differences are often absent between autistic males and their typically developing counterparts (Lemon et al., 2011; Nydén et al., 2000). Females with ASD also exhibit more widespread cognitive deficits and greater neuroanatomical abnormalities than males (Bloss & Courchesne, 2007; Rinehart et al., 2011). These findings suggest that if an inhibitory gating deficit exists in autistic traits, it is likely to be more prominent in females. However, no study to date has examined sex differences in working memory gating using the AA paradigm.

Beyond sex differences, the social versus non-social nature of stimuli may also play a role, particularly in females. Social camouflaging—defined as the use of effortful compensatory strategies to present a less visibly autistic persona during social interactions—is widely recognised as a key feature of the female autism phenotype (Hull et al., 2017; Lai et al., 2011). Autistic females consistently report higher levels of camouflaging behaviour than autistic males, reflecting greater engagement of top-down regulatory strategies to conform to social norms (Hull et al., 2020a). At its core, camouflaging relies on active inhibitory control over socially relevant information and responses. For high autistic trait females, this well-practised, domain-specific inhibitory mechanism may compensate for a general gating deficit when processing social stimuli, effectively masking the underlying impairment. For non-social stimuli that do not trigger camouflaging, by contrast, the general gating deficit would remain observable.

The present study employed the AA paradigm to investigate whether inhibitory working memory gating is reduced in individuals with high autistic traits, and whether this potential deficit is modulated by sex and stimulus type. In Experiment 1, we used non-social animal drawings as targets and measured surprise memory for task-irrelevant stimulus identity. In Experiment 2, we replaced targets with social face stimuli to test whether the gating pattern differs for social information. We predicted that high autistic trait females would show reduced inhibitory gating (higher surprise trial accuracy) for non-social stimuli relative to all other groups, but that this deficit would be eliminated for social stimuli due to the engagement of social camouflaging strategies.

**Experiment 1: Reduced inhibitory gating for non-social stimuli in high autistic trait females**

## Method

### Sample size

We conducted an a priori power analysis using G*Power 3.1 (Faul et al., 2007) to determine the required sample size for all experiments. Based on the effect size of the attribute amnesia effect (φ = 0.50) reported in Chen and Wyble (2015a), the analysis indicated that a minimum of 20 participants per group would be sufficient to achieve 80% power at an alpha level of 0.05. To ensure greater stability of the estimates, we set the sample size at 30 participants per group. Participants were not permitted to complete more than one experiment.

### Participants

Prior to the formal experiment, all participants completed the Beck Depression Inventory-I (BDI-I) and the Chinese Shorten Version of the Autism-Spectrum Quotient (the 15-item AQ-CSV). Participant selection followed the screening criteria established by Mahmoud (2023) and Xu et al. (2025). Individuals with a BDI-I score below 11 and an AQ score of 39 or above were assigned to the high autistic trait (HAT) group, whereas those with a BDI-I score below 11 and an AQ score below 39 were assigned to the low autistic trait (LAT) group. A total of 80 undergraduate students (40 females and 40 males) were recruited, with 40 participants (20 females and 20 males) in each group. Participants ranged in age from 18 to 25 years. All participants had normal or corrected-to-normal vision, were right-handed, and had no prior experience with similar experiments. No participant completed more than one experiment in this study. This study was approved by the local review board for the ethical treatment of human participants.

### Apparatus and Stimuli

All visual stimuli were selected from the standardized set of black-and-white line drawings developed by Snodgrass and Vanderwart (1980). The final stimulus set included 12 object drawings and 4 animal drawings, all of which were presented to each participant across the experiment. Each image was displayed at a resolution of 300 × 300 pixels, and the assignment of stimuli to individual trials was fully randomized.

Stimuli were presented on a 14-inch Lenovo monitor with a 60 Hz refresh rate and a native resolution of 2880 × 1800 pixels. The experiment was programmed in MATLAB (The MathWorks, Natick, MA) using the Psychophysics Toolbox extensions (Brainard, 1997; Pelli, 1997). Participants were seated with their head stabilized by a chin rest at a viewing distance of 50 cm from the screen, and made all responses via a standard computer keyboard.

### Design and paradigm

The experimental protocol was adapted from the classic attribute amnesia (AA) paradigm (Chen & Wyble, 2015a, 2015b, 2016; Chen et al., 2019). The trial sequence is illustrated in Figure 1. Each standard trial consisted of four phases: (1) a fixation phase, in which a black central fixation cross (0.62° of visual angle) and four black placeholder circles (0.62° × 0.62°) were displayed at the four corners of an invisible 6.25° × 6.25° square centered on the screen, with fixation duration varying randomly between 800 ms and 1800 ms; (2) a stimulus presentation phase, in which a stimulus array containing one target animal picture and three distractor object pictures was presented for 1000 ms, with each picture occupying one of the four placeholder locations; (3) a response phase, in which a 100-ms visual mask was presented following stimulus offset, followed by a 500-ms central fixation cross, after which participants completed an unspeeded spatial location judgment: four black digits (1–4) appeared at the four placeholder positions, and participants pressed the corresponding number key to report the location of the target animal, with the digits remaining on screen until a response was registered; and (4) a feedback phase, in which accuracy feedback was presented for 500 ms immediately after each response.

**Figure 1.**
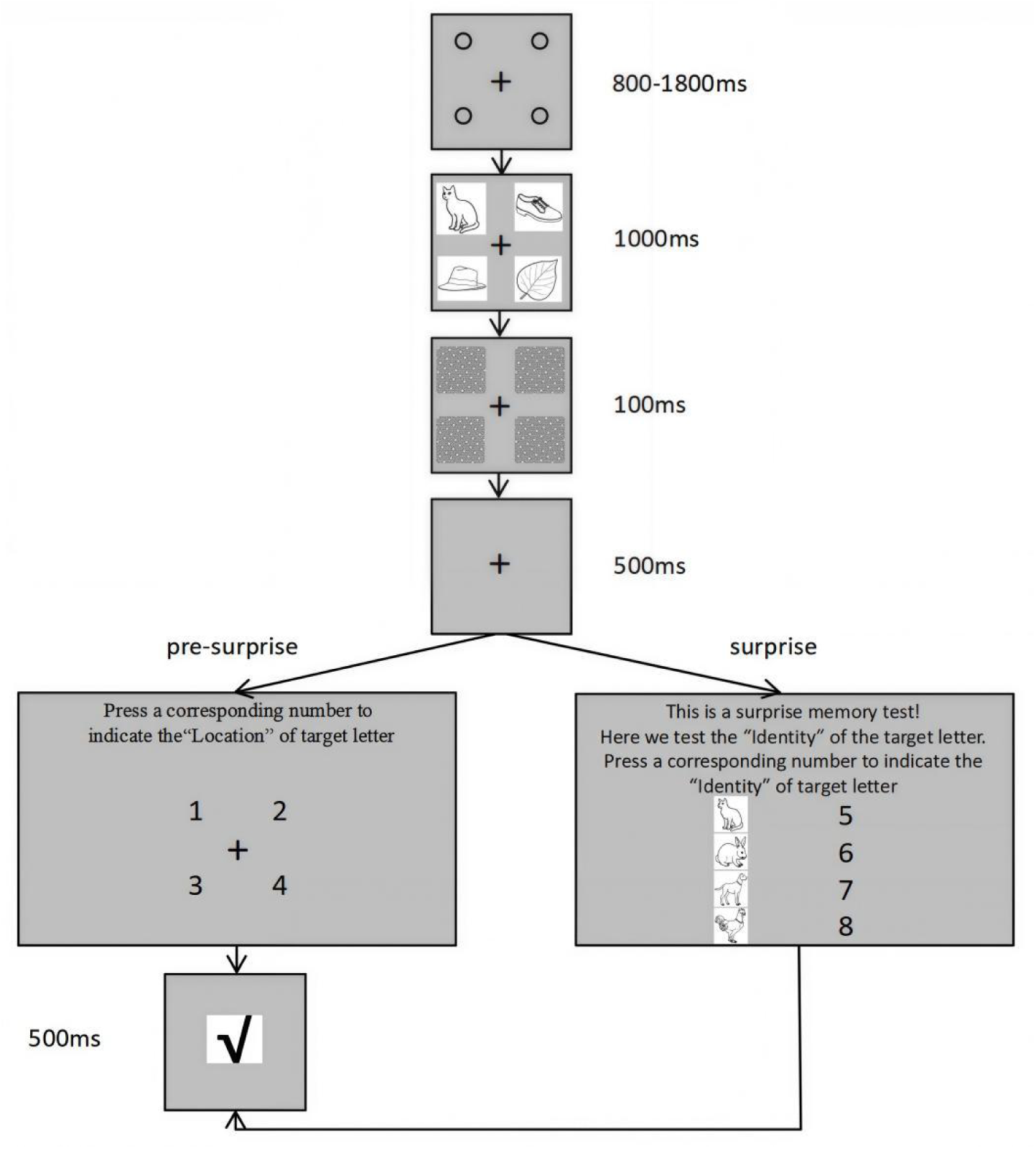
Schematic representation of the trial sequence for Experiment 1. In pre-surprise trials, participants performed a spatial localization task, indicating the position of the target animal image among three object distractor drawings. In the critical surprise trial, participants were unexpectedly instructed to first judge the identity of the target animal prior to making the standard location response. Subsequent control trials adopted the identical trial structure as the surprise trial. Stimulus dimensions are for illustrative purposes only and not drawn to scale.

Participants completed 160 trials in total. The first 155 trials followed the standard trial structure described above. The 156th trial served as a surprise trial, which incorporated an additional forced-choice target identification task. In this task, four animal pictures were presented simultaneously, and participants were instructed to select the target animal that had appeared in the preceding array. Four digits (5–8) were placed to the right of each picture, and participants pressed the corresponding key to indicate their choice; the stimuli remained visible until a response was made.

The presentation order of the identity discrimination task was counterbalanced across participants, and the spatial positions of the four animal pictures were randomized. The surprise trial concluded with the standard location judgment component. Following the surprise trial, participants completed four control trials, which were identical in task structure and trial sequence to the surprise trial.

## Results

As shown in Table 1, in the surprise trial, 9 out of 20 (45%) females in the low autistic trait group and 7 out of 20 (35%) males in the low autistic trait group correctly reported the animal identity, while performance on the location judgment task remained high across both groups. As illustrated in Figure 2, performance on the identity judgment task significantly improved in the first control trial for both groups: for females, from 45% to 95%, χ² (1, *N* = 40) = 8.10, *p* = 0.002, *φ* = 0.45; for males, from 35% to 80%, χ² (1, *N* = 40) = 7.11, *p* = 0.004, *φ* = 0.42.

**Figure 2.**
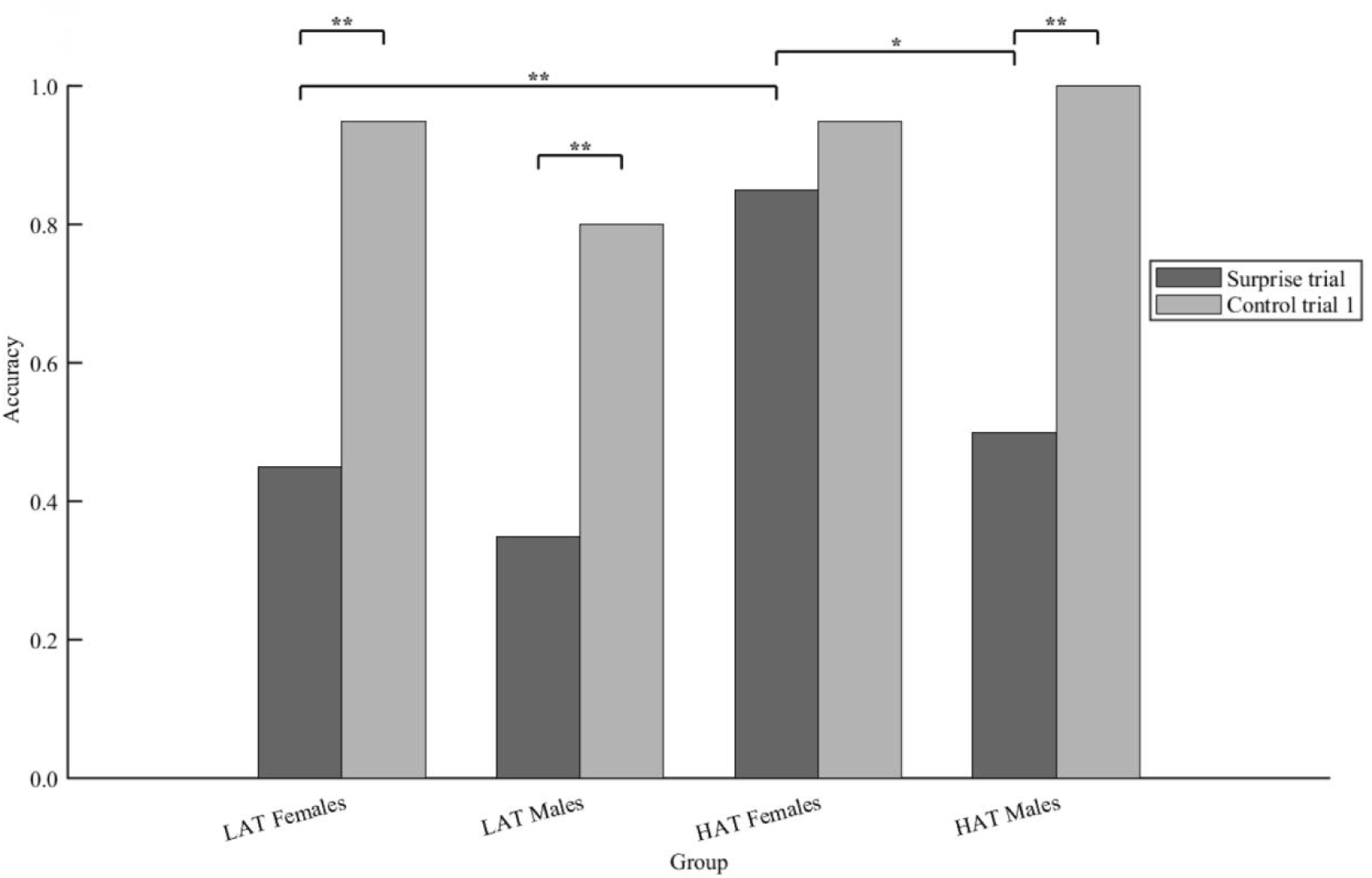
Animal Identity report accuracy in Experiment 1 as a function of group and trial type. Accuracy on the Surprise trial and Control trial 1 are shown separately for each group. *p <.05. **p <.01. ***p <.001. LAT = low autistic traits; HAT = high autistic traits. Control trial 1 = the first control trial in which participants were aware that identity judgment would be required.

**Table 1.** Accuracy in Experiment 1 (20 participants in each group)

| Task | Group | Sex | Pre-surprise | Surprise | Control 1 | Control 2 | Control 3 | Control 4 |
| --- | --- | --- | --- | --- | --- | --- | --- | --- |
| Location | low autistic trait | male | 100.00% | 70.00% | 85.00% | 95.00% | 95.00% | 95.00% |
|  |  | female | 100.00% | 75.00% | 100.00% | 100.00% | 100.00% | 100.00% |
|  | high autistic trait | male | 100.00% | 65.00% | 100.00% | 100.00% | 100.00% | 100.00% |
|  |  | female | 100.00% | 85.00% | 95.00% | 100.00% | 95.00% | 100.00% |
| Identity | low autistic trait | male | N/A | 35.00% | 80.00% | 90.00% | 100.00% | 100.00% |
|  |  | female | N/A | 45.00% | 95.00% | 100.00% | 100.00% | 100.00% |
|  | high autistic trait | male | N/A | 50.00% | 100.00% | 100.00% | 100.00% | 100.00% |
|  |  | female | N/A | 85.00% | 95.00% | 100.00% | 100.00% | 95.00% |

Note. N/A = not available. Controls 1-4 correspond to the four control trials.

In the surprise trial, 17 out of 20 (85%) females in the high autistic trait group and 10 out of 20 (50%) males in the high autistic trait group correctly reported the animal identity, whereas accuracy on the location judgment task remained high for both. In the first control trial, performance on the identity judgment task did not significantly change for high autistic trait females (from 85% to 95%), χ² (1, *N* = 40) = 0.25, *p* = 0.625, *φ* = 0.08. In contrast, high autistic trait males showed a significant improvement (from 50% to 100%), χ² (1, *N* = 40) = 8.10, *p* = 0.002, *φ* = 0.45. These patterns indicate that the relatively poor identity judgment accuracy observed in low autistic trait females, low autistic trait males, and high autistic trait males was not attributable to the unexpected nature of the surprise trial per se, but rather reflected a failure to encode the identity attribute. By contrast, high autistic trait females maintained high accuracy on the identity judgment task, demonstrating successful encoding of the identity attribute even when it was not explicitly required.

Between-group comparisons in the surprise trial revealed a significant difference between low and high autistic trait females, χ² (1, *N* = 40) = 7.03, *p* = 0.008, *φ* = 0.42, and no significant difference between low and high autistic trait males, χ² (1, *N* = 40) = 0.92, *p* =.337, *φ* = 0.15. Within the high autistic trait group, females significantly outperformed males, χ² (1, *N* = 40) = 5.58, *p* =.018, *φ* = 0.37. These results indicate a sex difference in the ability to consciously perceive the task-irrelevant attribute among individuals with high autistic traits. Performance on both the location and identity judgment tasks remained stable across the final three control trials.

### Experiment 2: Camouflaging of gating deficits for social stimuli in high autistic trait females Method

Experiment 2 followed the same procedure as Experiment 1, with one modification: the target stimuli were faces rather than animal drawings. A new sample of 160 participants was recruited for this experiment. Participants were divided into four groups based on a 2 (autistic trait level: high vs. low) × 2 (target identity: self vs. other) between-subjects design, with 20 males and 20 females in each cell (40 participants per condition). Thus, in the high autistic trait group, 40 participants were assigned to the self-face condition and 40 to the other-face condition; the low autistic trait group followed the same allocation.

The stimulus set consisted of the same 12 object line drawings used in Experiment 1, plus four face images per participant: one self-face photograph and three sex-matched unfamiliar faces. All images were cropped into circular shapes, resized to 300 × 300 pixels, and converted to grayscale. To minimise low-level visual differences, we applied the SHINE toolbox (Willenbockel et al., 2010) to equate luminance and contrast across all images (see Figure 3 for example stimuli).

**Figure 3.**
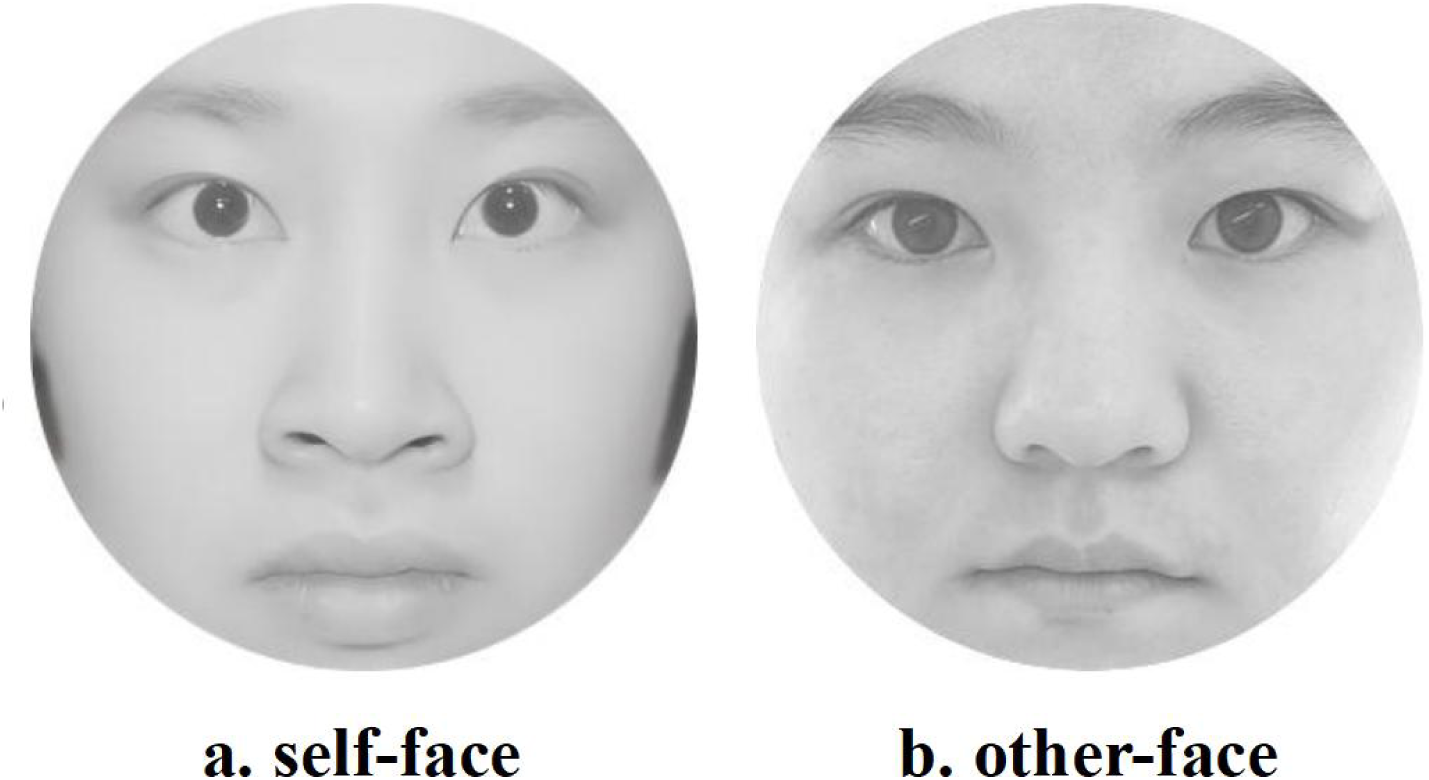
Example stimuli used in Experiment 2. (a) Self-face stimuli. (b) Other-face stimuli. Both face images are from the authors.

The trial sequence was identical to that of Experiment 1. In the first 155 trials, participants performed a location judgment task, indicating where the target face appeared among three distractor object drawings. On the 156th trial (surprise trial), participants were first asked to identify which face they had just seen, prior to reporting its location. This was followed by four control trials, which were identical in structure to the surprise trial (i.e., both identity and location judgments were required).

## Results

As shown in Table 2 and Figures 4–5, both the high and low autistic trait groups exhibited a robust attribute amnesia effect, as evidenced by significantly lower accuracy on the surprise trial than on the second control trial, as indicated by McNemar tests (all *p* < 0.039). Importantly, identity accuracy on the remaining three control trials (i.e., the second, third, and fourth) was consistently high across all conditions, reaching near-ceiling levels and confirming that the attribute amnesia effect was confined to the surprise trial.

**Figure 4.**
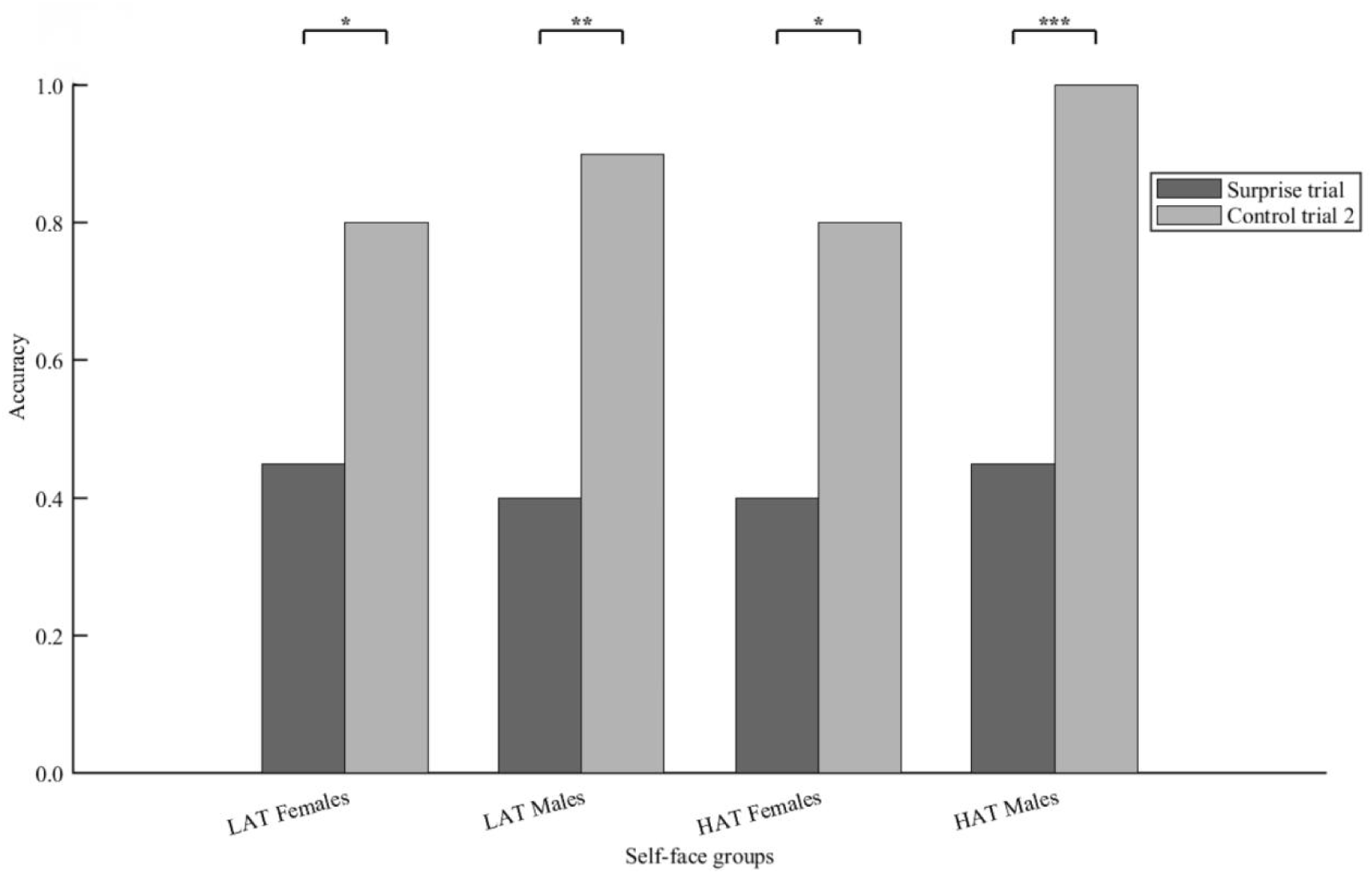
Face Identity report accuracy in Experiment 2 as a function of self-face group and trial type. Accuracy on the Surprise trial and Control trial 2 are shown separately for each group. *p <.05. **p <.01. ***p <.001. LAT = low autistic traits; HAT = high autistic traits. Control trial 2 = the second control trial in which participants were aware that identity judgment would be required.

**Figure 5.**
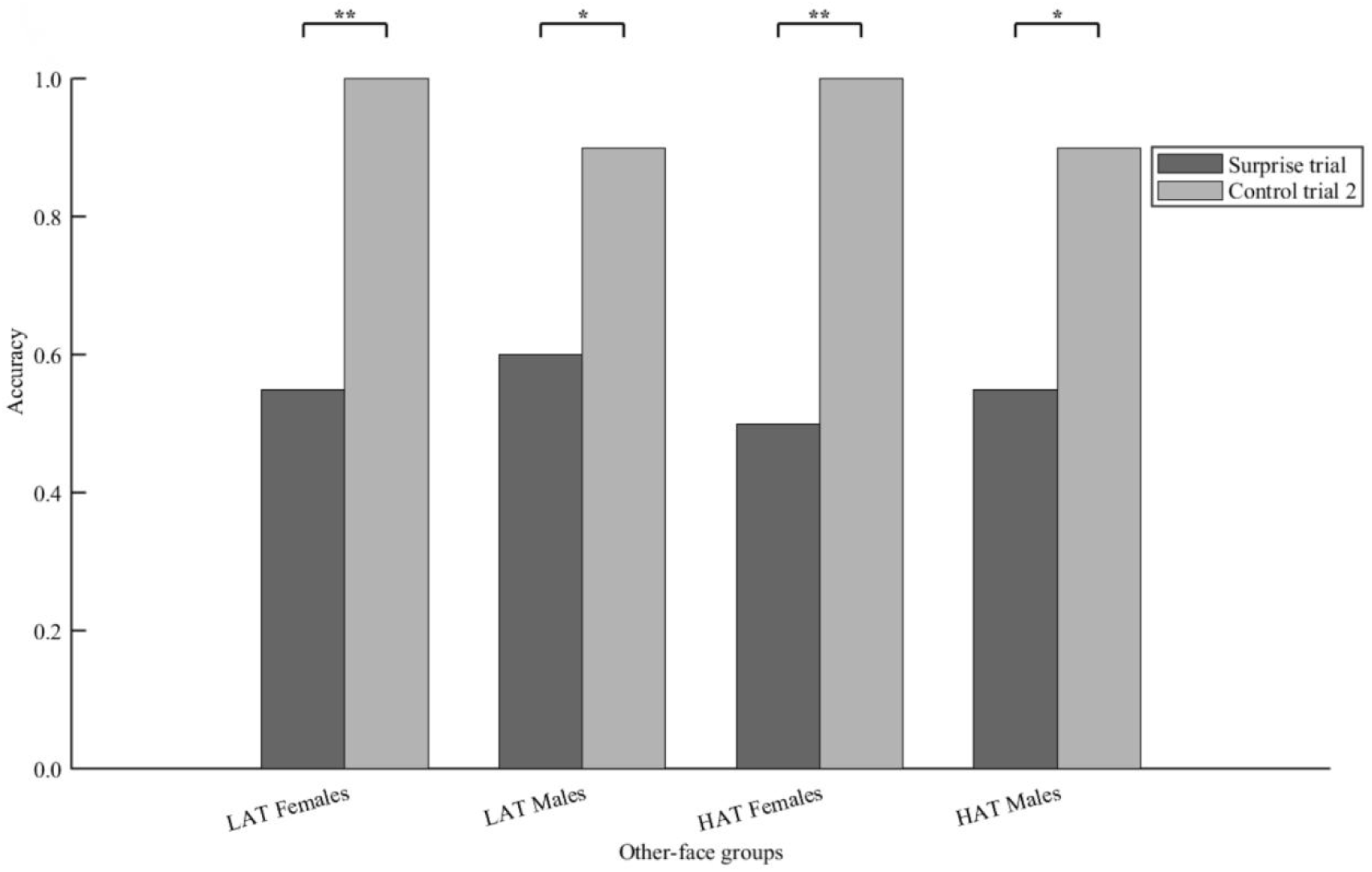
Face Identity report accuracy in Experiment 2 as a function of other-face group and trial type. Accuracy on the Surprise trial and Control trial 2 are shown separately for each group. *p <.05. **p <.01. ***p <.001. LAT = low autistic traits; HAT = high autistic traits. Control trial 2 = the second control trial in which participants were aware that identity judgment would be required..

**Table 2.**
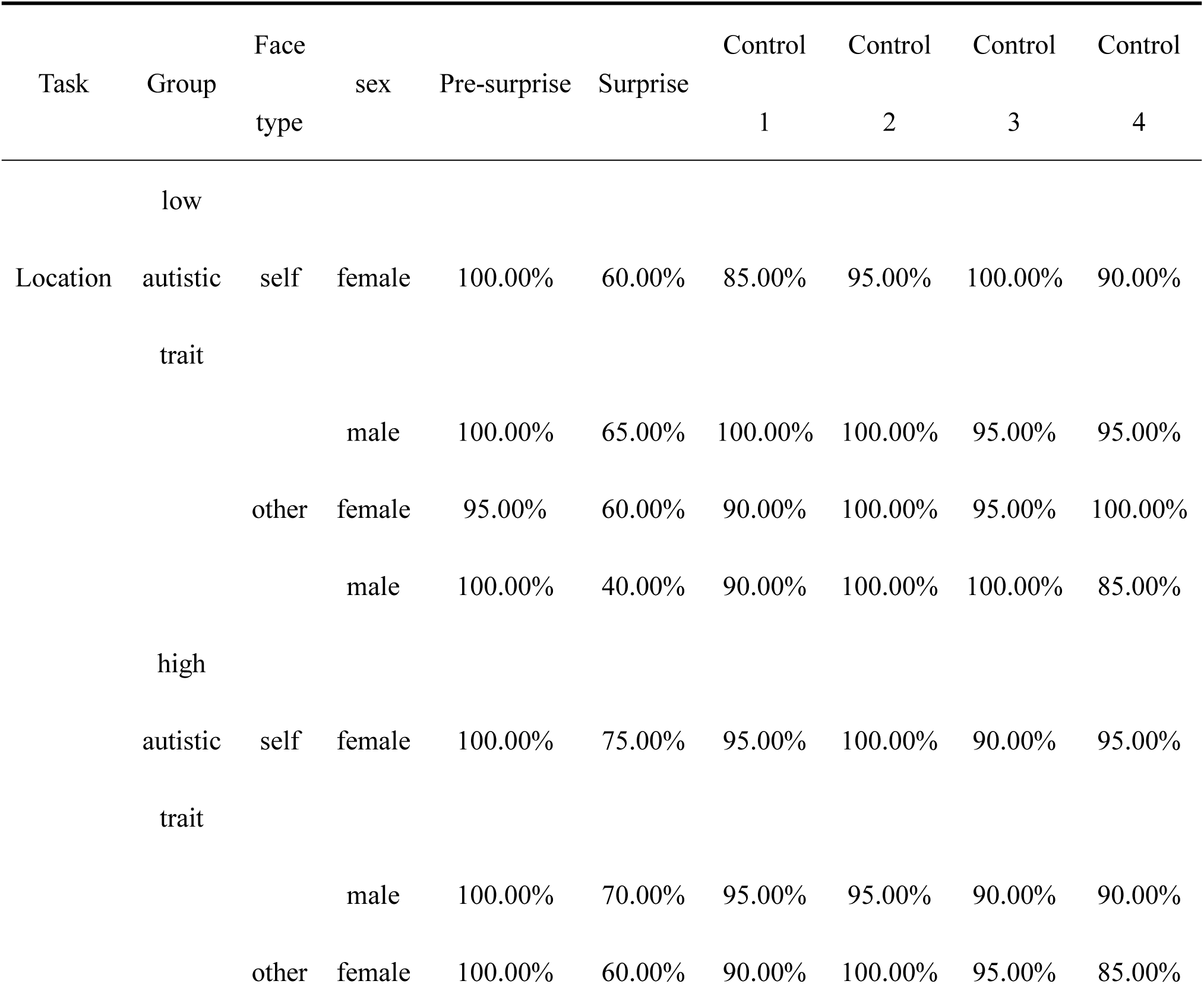
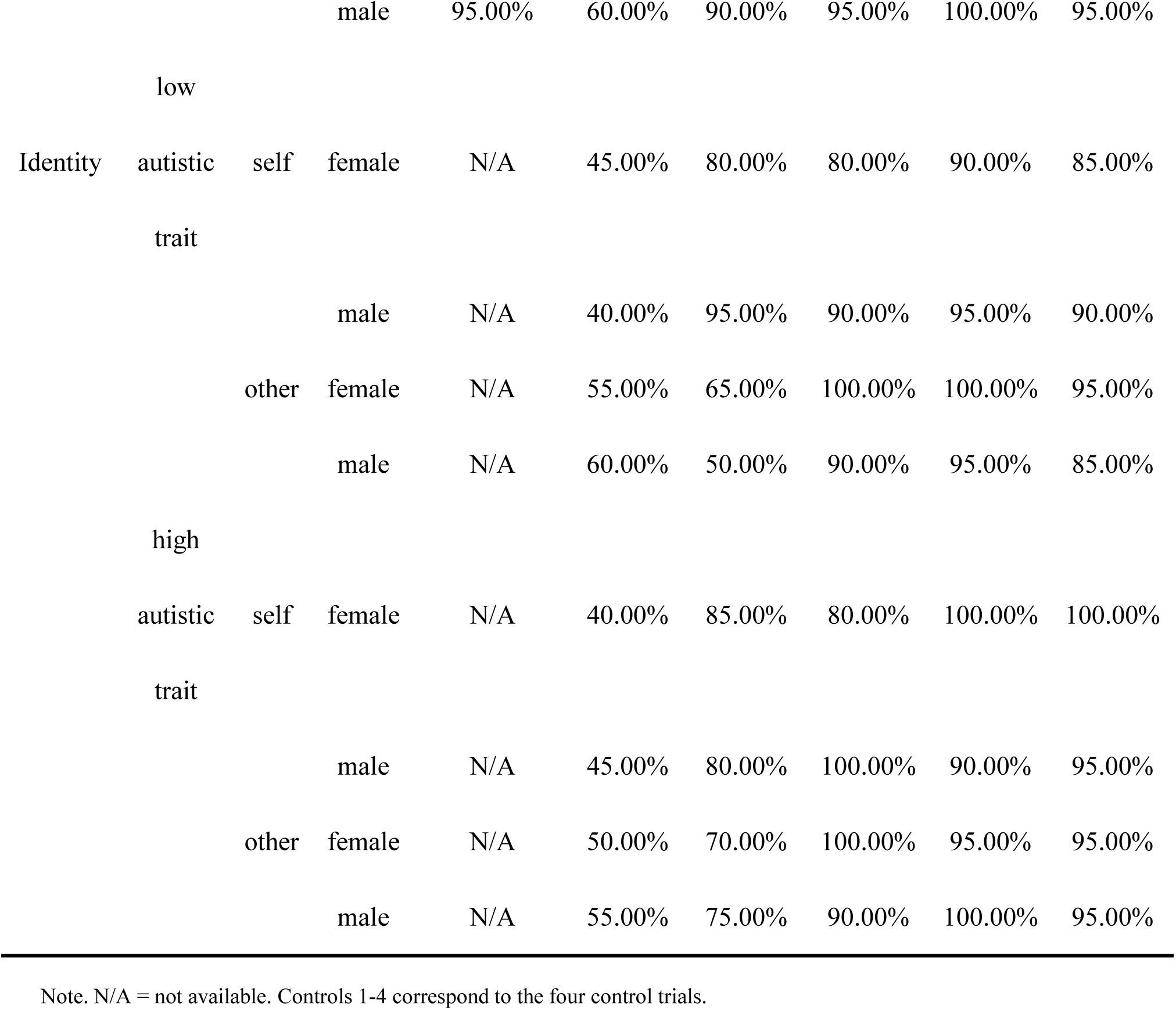
Accuracy in Experiment 2 (20 participants in each group)

To further examine whether autistic traits, sex, and face type modulated identity memory on the surprise trial, we conducted a binary logistic regression with identity accuracy (0 = incorrect, 1 = correct) as the dependent variable. The predictors included the three main effects (autistic trait group: high vs. low; sex: male vs. female; face type: self vs. other), the three two-way interaction terms (autistic trait group × sex, autistic trait group × face type, sex × face type), and the three-way interaction term (autistic trait group × sex × face type). All predictors were entered simultaneously using the Enter method. The main effects of autistic trait group (Wald χ² = 0.102, *p* = 0.749), sex (Wald χ² = 0.102, *p* = 0.749), and face type (Wald χ² = 0.403, *p* = 0.526) were all non-significant. None of the interaction terms reached significance: autistic trait group × sex (*p* = 0.651), autistic trait group × face type (*p* = 0.996), sex × face type (*p* = 0.996), or the three-way interaction autistic trait group × sex × face type (*p* = 0.746). The overall model was not significant, χ² (7) = 3.118, *p* = 0.874, Nagelkerke R^2^ = 0.026, indicating that none of the predictors reliably distinguished identity memory performance. These results suggest that when the task-irrelevant attribute is social in nature, the gating deficit observed for non-social stimuli is no longer expressed, with all groups performing near chance levels on the surprise identity judgment.

To quantify the evidence for the absence of group differences, we conducted Bayesian contingency table analyses comparing high versus low autistic trait females separately for self-face and other-face conditions. For self-faces, the Bayes factor (BF_01_) was 2.565, indicating that the data were 2.57 times more likely under the null hypothesis (no difference between groups) than under the alternative hypothesis. For other-faces, BF_01_ was 2.542, providing comparable evidence for the absence of a group difference. The log odds ratios for both comparisons had 95% credible intervals that crossed zero (self-face: [-1.042, 1.408]; other-face: [-1.018, 1.383]), further confirming that high autistic trait females did not outperform low autistic trait females in either face condition. These findings provide weak-to-moderate anecdotal evidence that the inhibitory gating deficit in high autistic trait females is masked when processing social stimuli, regardless of whether the face is self or other.

### General discussion

The present study employed the attribute amnesia paradigm to investigate whether inhibitory working memory gating is reduced in individuals with high autistic traits, and whether this potential deficit is modulated by stimulus type and sex. Across two experiments, we found a clear dissociation: when the task-irrelevant attribute was non-social (animal identity), high autistic trait females showed significantly higher accuracy on the surprise trial than all other groups, indicating reduced active inhibitory gating efficiency. However, when the task-irrelevant attribute was social (face identity), this gating advantage disappeared entirely, as all groups performed at approximately chance levels on the surprise trial despite performing well on control trials. These findings suggest that the inhibitory gating deficit in high autistic trait females is not a global impairment, but rather one that is modulated by the social versus non-social nature of the stimulus. We interpret this pattern within a dual-mechanism framework: for non-social stimuli, the deficit in general-purpose active inhibitory gating is exposed; for social stimuli, a domain-specific compensatory strategy (i.e., social camouflaging) is recruited in females, which imposes extra top-down attentional control over task-irrelevant social attributes and effectively masks the expression of the underlying gating deficit.

### Gating deficits in high autistic trait females for non-social stimuli **(animal identity)**

In Experiment 1, high autistic trait females correctly reported the animal identity on the surprise trial at substantially higher rates (85%) than all other groups (45%, 35%, and 50% for low autistic trait females, low autistic trait males, and high autistic trait males, respectively), while all groups performed accurately on the location judgment task. Crucially, all groups improved significantly on the first control trial, confirming that identity information had been successfully encoded. The selective preservation of identity information in high autistic trait females therefore reflects a failure to actively inhibit task-irrelevant information from entering working memory, rather than a failure to encode that information (Chen & Wyble, 2015a; Liu et al., 2025).

Drawing on the weak central coherence account (Shah & Frith, 1993; Happé & Frith, 2006) and the enhanced perceptual functioning model (Mottron et al., 2006), this gating deficit could be understood as arising from the conjunction of enhanced local processing and weakened top-down inhibitory control in high autistic trait females. When enhanced perceptual encoding of task-irrelevant information combines with reduced inhibitory efficiency, this information is automatically maintained in working memory and becomes accessible on the surprise trial.

The sex specificity of this effect is noteworthy. High autistic trait males performed similarly to low autistic trait groups. The observed sex difference is broadly consistent with prior evidence that autistic females, compared to autistic males, carry a greater load of genetic variants (Gilman et al., 2011; Levy et al., 2011) and show more pronounced neurodevelopmental differences (Bloss & Courchesne, 2007; Schumann et al., 2010). At the behavioural level, this pattern aligns with evidence that females with ASD or high autistic traits exhibit more pronounced impairments than males in response inhibition, cognitive flexibility, and working memory (Lemon et al., 2011; Nydén et al., 2000; Memari et al., 2013; Kiep & Spek, 2017; Rinehart et al., 2011). The deficit observed in high autistic trait females thus emerges from the conjunction of high autistic traits with female sex. Together, these findings demonstrate that high autistic trait females exhibit reduced inhibitory gating efficiency at the attentional input stage, manifested as insufficient filtering of task-irrelevant non-social perceptual attributes, and that this deficit is sex-specific.

### Absence of Gating Deficits in High Autistic Trait Females for Social Stimuli

In Experiment 2, high autistic trait females did not show preserved processing of face identity information on the surprise trial. Instead, all groups, regardless of autistic trait level or sex, performed at approximately 50% accuracy on the identity judgment task, while performing accurately on the location task. Crucially, all groups improved substantially on the control trials, confirming that identity information had been successfully encoded and was retrievable when explicitly required. This stands in stark contrast to the pattern observed in Experiment 1, where high autistic trait females showed preserved processing of task-irrelevant non-social information (animal identity). Therefore, the absence of a gating advantage in Experiment 2 cannot be attributed to a failure to encode identity information. Rather, it suggests that when the task-irrelevant attribute is social in nature, the gating deficit observed for non-social stimuli is no longer expressed.

We propose that this reflects the engagement of social camouflaging, defined as the use of strategies to present a less visibly autistic persona during social interactions (Hull et al., 2017). Camouflaging, like autistic traits, is thought to exist on a continuum across the entire population (Hull et al., 2020a). Camouflaging is fundamentally an inhibitory process that requires the suppression of visible or detectable autistic characteristics through sustained top-down attentional control (Johnson et al., 2015; Livingston & Happé, 2017). This inhibitory nature maps directly onto the cognitive demands of the attribute amnesia paradigm: to prevent task-irrelevant identity information from being reported on the surprise trial, participants must actively suppress that information from entering or remaining in working memory. For high autistic trait females, who have extensive experience deploying camouflaging strategies in daily social contexts (Bargiela et al., 2016), this inhibitory process may be particularly well-practised and readily triggered by social stimuli.

Consistent with this interpretation, meta-analytic evidence confirms that autistic females consistently score higher than autistic males across all camouflaging dimensions (Hull et al., 2020a; Canciño-Barros et al., 2025). This sex difference emerges early in development: autistic girls show greater social reciprocity than autistic boys despite similar levels of autistic traits (Wood-Downie et al., 2021). Importantly, this pattern appears to be specific to autistic individuals; non-autistic males and females do not differ in their camouflaging behaviours (Dean et al., 2017; Hull et al., 2020a). Camouflaging has been argued to be a key feature of the female autism phenotype (Hull et al., 2020a; Wood-Downie & Wong, 2017), and autistic females show higher levels of compensatory camouflaging (Lai et al., 2017), supported by superior executive function abilities that underpin sustained top-down social regulation (Lehnhardt et al., 2016), consistent with the proposition of “improved behavioural presentation despite persisting core deficit(s)” (Livingston & Happé, 2017). This may be partly driven by a stronger tendency to systemise social behaviour in autistic females compared to males (Hull et al., 2017). Notably, camouflaging is often less frequent in the presence of close friends and family (Hull et al., 2017), whereas the faces used in the present study were all unfamiliar others, a context likely to elicit greater camouflaging.

Taken together, the cross-experiment dissociation supports a dual-mechanism framework of inhibitory gating in high autistic trait females: a general inhibitory gating deficit is evident for non-social stimuli, but is behaviourally masked by domain-specific social camouflaging during social processing. This stimulus-dependent pattern demonstrates that the working memory gating impairment in high autistic trait females is not a global deficit, but is modulated by stimulus sociality and the recruitment of compensatory strategies.

### Limitation

One additional observation warrants consideration. In Experiment 2, location accuracy on the surprise trial was lower relative to pre-surprise and control trials. This likely reflects the fact that the location judgment was administered after the identity question, allowing decay or interference to disrupt spatial information (Chen & Wyble, 2016). Moreover, after excluding outliers (Tukey’s fences method, Q3 + 1.5 × IQR), identity judgment took significantly longer in Experiment 2 (*M* = 7010.29 ms, *SD* = 3392.91) than in Experiment 1 (*M* = 4807.07 ms, *SD* = 3092.07), *t*(222) =-4.686, *p* < 0.001, suggesting that longer processing time for faces may have allowed more decay of location information. The holistic nature of face processing (Tanaka & Farah, 1993; Robbins & McKone, 2007) may also impose greater cognitive load, interfering with spatial maintenance (Langton et al., 2008). These factors together account for the greater location accuracy drop observed in Experiment 2.

Regarding identity performance, self-face accuracy on Control Trial 1 recovered to near-ceiling levels (80%, 95%, 85%, and 80% across groups), whereas other-face accuracy did not (65%, 50%, 70%, and 75%). This asymmetry suggests that self-faces enjoy an encoding advantage that can be realised once reporting expectations are established (Alzueta et al., 2019). In the absence of such expectations (on the surprise trial), the gating mechanism remains closed (Miller & Cohen, 2001; McNab & Klingberg, 2008), and self-face identity cannot enter working memory. Other-face identities, lacking this advantage, show slower recovery. Critically, accuracy on Control Trials 2–4 was consistently high for both self-face and other-face conditions, confirming that participants were fully capable of encoding identity once expectations were established (Chen & Wyble, 2015a, 2016). The modest other-face performance on Control Trial 1 likely reflects the additional cognitive demands of reconfiguring encoding strategies immediately following the unexpected surprise probe (Chen et al., 2019), rather than a fundamental encoding deficit.

## Conclusion

The present study employed the attribute amnesia paradigm to investigate inhibitory working memory gating in individuals with high and low autistic traits, with particular attention to sex differences and stimulus specificity. Across two experiments, we found that high autistic trait females showed reduced gating of task-irrelevant non-social information (animal identity), as evidenced by higher accuracy on a surprise memory test. However, this gating advantage was absent when the task-irrelevant information was social (face identity), a finding we interpret as reflecting the engagement of social camouflaging, an inhibitory strategy that is particularly developed in females. These findings demonstrate that the inhibitory gating deficit in high autistic trait females is not a global impairment, but rather a stimulus-dependent one: non-social stimuli expose the deficit, whereas social stimuli recruit domain-specific camouflaging strategies that mask it. This dual-mechanism framework contributes to understanding how autistic traits manifest differently across sexes and stimulus domains, and underscores the importance of considering sex differences and stimulus characteristics in future research on autistic trait cognitive mechanisms.

## Author Note

This research was supported by the National Key Research and Development Program of China (2025YFE0213500), the National Science Foundation of China (32371098), Guangdong Basic and Applied Basic Research Foundation (2024A1515011429) and the Research Center for Brain Cognition and Human Development, Guangdong, China (No. 2024B0303390003).

We have no conflicts of interest to disclose.

## Notes

### Competing Interest Statement

The authors have declared no competing interest.

